# Harnessing patient-specific response dynamics to optimize evolutionary therapies for metastatic clear cell renal cell carcinoma – Learning to adapt

**DOI:** 10.1101/563130

**Authors:** I. C. Sorribes, A. Basu, R. Brady, P. M. Enriquez-Navas, X. Feng, J. N. Kather, N. Nerlakanti, R. Stephens, M. Strobl, I. Tavassoly, N. Vitos, D. Lemanne, B. Manley, C. O’Farrelly, H. Enderling

## Abstract

Renal cell carcinoma (RCC) is one of the ten most common and lethal cancers in the United States. Tumor heterogeneity and development of resistance to treatment suggest that patient-specific evolutionary therapies may hold the key to better patients prognosis. Mathematical models are a powerful tool to help develop such strategies; however, they depend on reliable biomarker information. In this paper, we present a dynamic model of tumor-immune interactions, as well as the treatment effect on tumor cells and the tumor-immune environment. We hypothesize that the neutrophil-to-lymphocyte ratio (NLR) is a powerful biomarker that can be used to predict an individual patient’s response to treatment. Using randomly sampled virtual patients, we show that the model recapitulates patient outcomes from clinical trials in RCC. Finally, we use *in silico* patient data to recreate realistic tumor behaviors and simulate various treatment strategies to find optimal treatments for each virtual patient.

## 1. Introduction

Renal cell carcinoma (RCC) represents a heterogeneous group of malignancies composed of a diverse histologic and genetic assembly [1, 2, 3]. It is one of the most common and lethal cancers in the United States. Clear cell renal cell carcinoma (ccRCC) is the most common subtype of RCC, presenting metastasis by the time of diagnosis in nearly one-third of patients [2]. The progression of ccRCC is modulated, in part, by immune cells in the tumor microenvironment. Cytotoxic T cells can prevent tumor growth, but tumor-associated suppressor cells, such as regulatory T cells, can suppress T cells response allowing the tumor to further develop [4].

For metastatic ccRCC (m-ccRCC) patients, cytotoxic chemotherapy offers little to no survival benefit [5]. The mutational phenotype of m-ccRCC patients suggests vascular endothelial growth factor (VEGF) and mechanistic target of rapamycin (mTOR) pathway as potential targets for successful treatment [6]. However, the use of targeted therapies, which include tyrosine kinase inhibitors (TKi) and mTOR inhibitors (mTORi), has extended the survival of patients with m-ccRCC only moderately, with an average of 14 to 24 months [7]. More recently, immunotherapeutics, including immune checkpoint inhibitors such as nivolumab (Nivo) and others have been approved by the FDA for m-ccRCC. Yet, m-ccRCC remains a lethal disease due to development of resistance [8]. None of the current therapies consider patient-and tumor-specific factors, and have yet to be optimized in their timing or sequencing [9, 3, 10].

Additionally, continuous treatment fails to consider the evolutionary dynamics of treatment response, where competition, adaptation, and selection between treatment sensitive and resistant cells contribute to therapy failure. In fact, continuous treatment maximally selects for resistant phenotypes and eliminates other competing populations, and may actually accelerate the emergence of resistant populations a well-studied evolutionary phenomenon termed competitive release [11, 12]. Intermittent therapies and alternating drug combinations have the potential to exploit these evolutionary dynamics to help extend patient survival [13]. However, identifying the specific treatment and treatment combinations that would benefit an individual patient is challenging [2].

Mathematical models that adequately simulate tumor growth, tumor-immune interactions, and treatment mechanisms may be calibrated to individual patients’ disease dynamics. Then, *in silico* simulations of parameterized models can test a wide variety of treatment strategies and ultimately find an optimal treatment strategy. Dynamic biomarkers may elucidate individual tumor sensitivity to different drugs to help develop evolutionary therapies [14]. Neutrophil-to-lymphocyte ratio (NLR), an inexpensive and frequently used blood test, may be a candidate for a biomarker of ccRCC response to treatment [15, 16, 17, 18]. We hypothesize that the NLR can inform the presented mathematical framework to help develop therapies that drive the evolution of the tumor-immune landscape towards control.

Our overall goal is to evaluate radiographic tumor responses and dynamic changes in the host tumor-immune landscape in sequentially obtained blood samples. These will serve to trigger treatment adaptations. We propose a system of three ordinary differential equations that account for tumor cells, effector immune cells, and tumor-associated suppressor immune cells. In our model, we account for the effect of three treatment strategies, namely mTORi, TKi, and Nivolumab. By simulating *in silico* responses to different therapies, we aim to identify previously-hidden functional sensitivity profiles. These profiles are used to simulate responses to the three drug classes on a per-patient basis *in silico* – at a magnitude not possible through traditional biological investigations alone.

## 2. Mathematical Model

We present a mathematical model of tumor-immune interactions based on the model proposed by Kuznetsov et al. [19]. Here we consider immune cells to be divided into two groups: effector cells that fight the tumor, and suppressor cells that inhibit effector cells, thereby benefiting tumor cells. Thus our model has three principal variables: *T*(*t*), the number of tumor cells at time *t*; *E*(*t*), the number of effector immune cells at time *t*; and *S* (*t*), the number of suppressor immune cells at time *t*, where time is measured in days. In this approach, we approximate the NLR with *S* (*t*)*/E*(*t*).

The increase in tumor cell numbers in the absence of treatment is assumed to follow logistic growth, with intrinsic growth rate *α* and carrying capacity *K*. They are killed by effector cells at rate *η*. Effector cells are assumed to be recruited to the tumor site. In our model, this process is divided into two events: normal influx of effector cells to the site, given by *σ*, and additional accumulation of cytotoxic cells in the region due to the presence of the tumor. Such additional accumulation was taking to follow the functional form proposed by Kuznetsov et al. [19], where *ρ* represents the maximum rate of accumulation, and *γ* the constant accumulation of immune cells at equilibrium. Effector cells die due to exhaustion at rate *µ*, and due to natural apoptosis at a rate *ω*. Suppressor cells are assumed to accumulate at the tumor site in response to the tumor, following the same functional form as effector cells. They inhibit the killing of tumor cells by effector cells and are also assumed to die at rate *ϵ*. Thus, without treatment, the model is given by

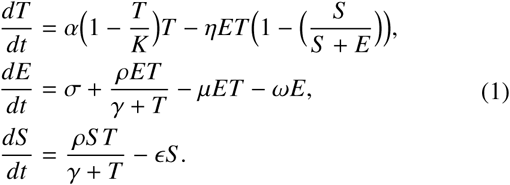

RCC treatments are classified based on their mechanisms of action: mTOR inhibitors, TK inhibitors, and immunotherapy. Thus we extend the system of equations in (1) to account for each treatment. mTORis are known to affect tumor cell proliferation and in our model they reduce *α* by rate *β*. TKis stop tumor cells from secreting VEGF, stopping angiogenesis and effectively starving tumor cells. In our model this is viewed as an increased reduction of the tumor growth rate for large cell numbers, represented by *φ*. Finally, Nivo increases the recruitment of effector cells by increasing the maximum rate of accumulation of effector cells by *ψ* percent. The values of *β, φ*, and *ψ* will determine the response or resistance of the tumor to each of the treatments. A schematic of the model is shown in Figure 1. Incorporating these treatments, the model becomes

**Figure 1:**
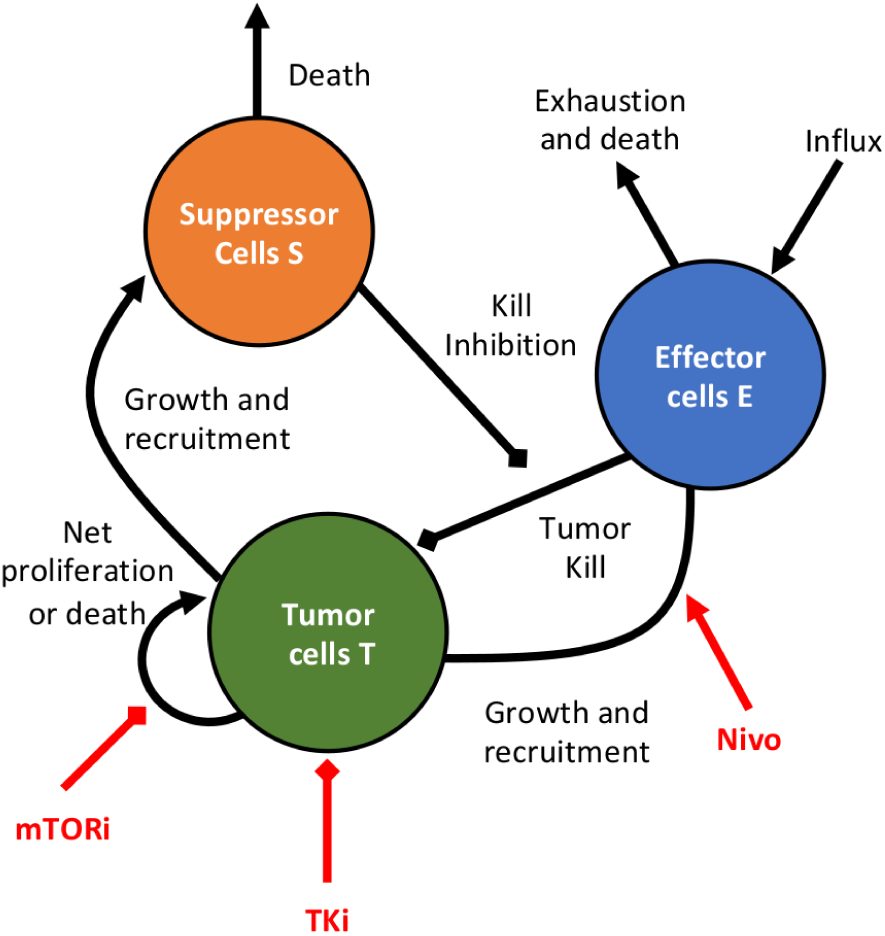
Model Schematic. The model is comprised of three variables: tumor *T*, effector *E*, and suppressor *S* cells. Tumor cells proliferate following a logistic growth and recruit immune cells. Effector cells have a constant influx to tumor site, induce tumor cell death, and can die due to exhaustion or apoptosis. Suppressor cells inhibit the killing of tumor cells by effector cells and can undergo apoptosis. mTORi treatments inhibit tumor growth, TKi causes starvation of tumor cells, and Nivo increases the recruitment of effector cells.

**Table 1:** Parameter values, units, and corresponding reference.

| Parameter | Value | Units | Reference |
| --- | --- | --- | --- |
| $\alpha$ | 0.18 | day<sup>-1</sup> | [19] |
| $K$ | $5 \times 10^9$ | cells | [19] |
| $\eta$ | $1.1 \times 10^{-8}$ | cells<sup>-1</sup> day<sup>-1</sup> | [19] |
| $\sigma$ | $1.3 \times 10^4$ | cells day<sup>-1</sup> | [19] |
| $\rho$ | 0.1245 | day<sup>-1</sup> | [19] |
| $\gamma$ | $2.019 \times 10^7$ | cells | [19] |
| $\mu$ | $3.422 \times 10^{-10}$ | cells<sup>-1</sup> day<sup>-1</sup> | [19] |
| $\omega$ | 0.0412 | day<sup>-1</sup> | [19] |
| $\epsilon$ | 0.0622 | day<sup>-1</sup> | Fixed |

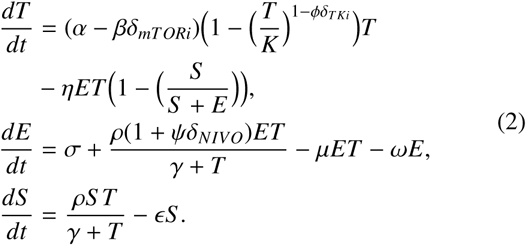

Each treatment has a Kronecker delta function *δ_mTORi_, δ_T Ki_, δ_NIVO_* that will be equal to one when treatment is being administered, and zero otherwise. All parameters in equations (1) where taken from Kuznetsov et al. [19], except for *ϵ*, which was fixed at a value 0.0622 (see Table 1). Thus our model is completely determined by the initial conditions of tumor, effector, and suppressor cells, and the resistance to each of the treatments *β, φ, ψ*. Initial conditions are assumed to be patient-specific and will determine patients’ prognoses.

### 2.1. Analysis

Given a specific choice of initial conditions and treatment parameters, we can determine tumor evolution by studying the dynamics of the system using phase plane analysis. First, consider the equations given in (1) of untreated tumor growth. We note that, since our model is used for biological purposes, we only consider positive cell populations, and positive parameters. Under these assumptions the nullclines for effector and tumor cells are given by

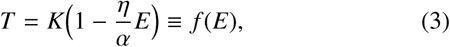

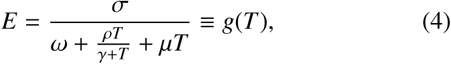

which are variations of the ones obtained by Kuznetsov et al. [19]. Furthermore, the steady states for the T-E phase plane are

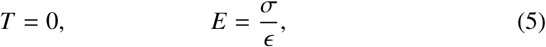

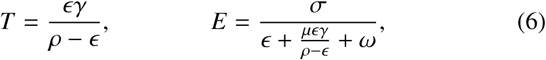

where (5) can be interpreted as tumor remission, and (6) is a saddle point which basins of attraction divides the phase plane into a responsive and non-responsive area of initial conditions. These steady-states show that without treatment even patients with a robust initial immune response will eventually exhaust effector cells allowing the tumor to escpape. Figure 2 shows the phase plane of effector versus tumor cells, along with their trajectories for varying initial conditions. Patients with a small tumor burden and relatively high effector cell population will show an initial equilibrium between tumor cells and the immune system, followed by inevitable tumor growth.

**Figure 2:**
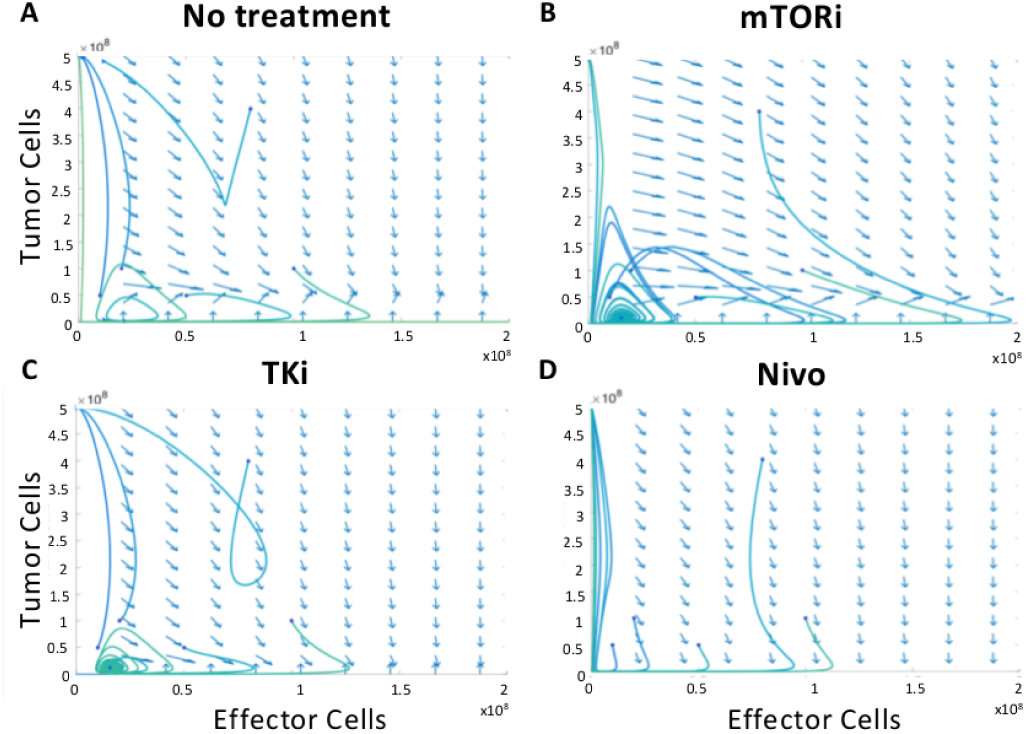
Phase planes under various treatment conditions. Phase planes of tumor versus effector cells considering **A.** no treatment and treatment after 20 days with **B.** mTORi, **C.** TKi, and **D.** Nivo **(D.)** Blue arrows represent the gradient field and solid curves are representative trajectories. Solid dots and arrows denote the beginning and end of the trajectory, respectively. Depending on the initial conditions, the tumor may enter remission.

However when treatment is applied, the saddle point and its basins of attraction move over time. Some non-responder patients can now move towards the remission steady state. As the treatment wears off or patients develop resistance, the trajectory shows two different behaviors. In one, the trajectory will cross towards the remission area before the basin of attraction returns to its original state, leading the patient to successfully reach remission. Alternatively, if the trajectory does not enter the remission area before the treatment’s effect disappears, the tumor will eventually regrow. In this case, patients exhibit a transient response.

Figures 2B, C, and D show the phase planes for mTORi, TKi, and Nivo therapy, respectively. Patients with the same initial conditions behave in different ways in each of the phase planes. Under mTORi, two patients will show an initial response but later develop resistance, while the remaining patients will converge to the remission state. With TKi, patients with higher tumor burden will not respond to treatment, while the others will show remission. Finally, all patients treated with Nivo show an initial response but treatment ultimately fails. Based on this knowledge, an optimal treatment should move the basin of attraction in such a way that the patients’ trajectory will be attracted to the remission steady-state long enough that the trajectory can cross to its area of attraction before treatment wears off.

## 3. Results

Patients with RCC can respond in different ways to the same treatment. Some patients show no response while others will show an initial response that can be lasting or temporary. From a mathematical point of view, we can correlate this notion with trajectories in our model. In Figure **??** we show trajectories for 25 virtual patients, created by randomly choosing initial conditions and treatment parameters (*β, φ, ψ*). In all cases, we applied Nivo after 30 days of tumor progression. Out of the 25 patients, ≈ 30% show a transient initial response, eventually developing resistance to treatment. The other virtual patients were not responsive to treatment.

### 3.1. NLR Stratifies Response

Figure 3A shows the Kaplan-Meier plots for our virtual patient cohort and demonstrates that patients with a low NLR have a significantly higher survival compared to those with a high NLR when Nivo is applied. These results are in good agreement with those by Bilen et al. who used a cutoff of 5.5 to stratify patients into NLR low or NLR high Bilen et al. [16]. Figure 3B shows the Kaplan-Meier plot for our group of virtual patients when mTORi and TKi are applied. We compared our results with those presented in Choueiri et al. [20] where, in an open-label randomized phase 3 trial, 658 patients with advanced or metastatic ccRCC were randomly assigned to receive cabozantinib (TKi) or everolimus (mTORi) once a day (see Figure 4 in [20]). Based on these results, we are confident that the structure of the model adequately recreated tumor-immune and treatment dynamics.

**Figure 3:**
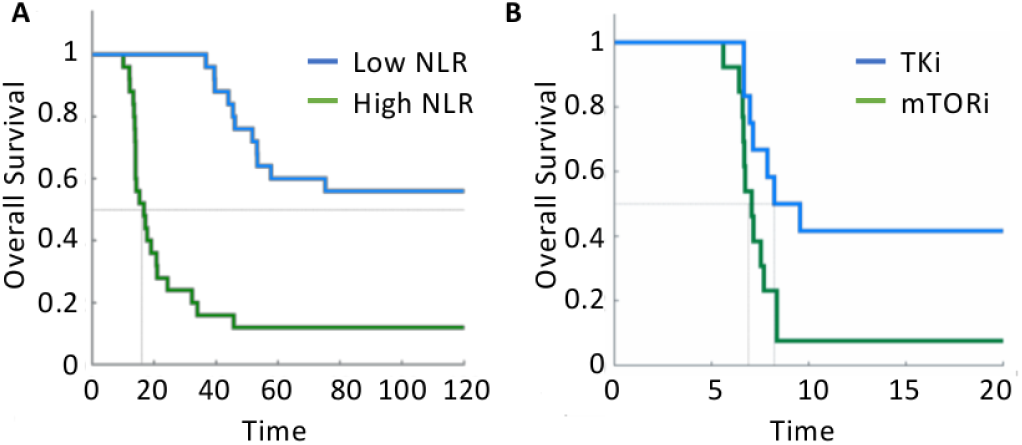
**A** Kaplan Meier plots of 25 virtual patients treated with Nivo, stratified by NLR (low NLR blue, high NLR green). **D** Kaplan Meier plots for virtual patients treated with TKi (blue) and mTORi (green).

### 3.2. Clinical Feasibility & Optimal Treatment Strategies

In order for our model to be clinically relevant, it must be able to identify patient-specific parameters with an acceptable degree of accuracy, with ideally as few time points as possible. To assess the number of data points required for acceptable accuracy, we performed *in silico* studies on our virtual patients for whom the true parameters are known. Figure 4, illustrates the results for one of our of our virtual patients, treated with immunotherapy. From this, we found that six weekly data points are sufficient to accurately capture the tumor behavior. With this conceptual exercise we show that, given one to two months of a patient’s data, we will be able to estimate the model parameters and accurately model tumor dynamics.

**Figure 4:**
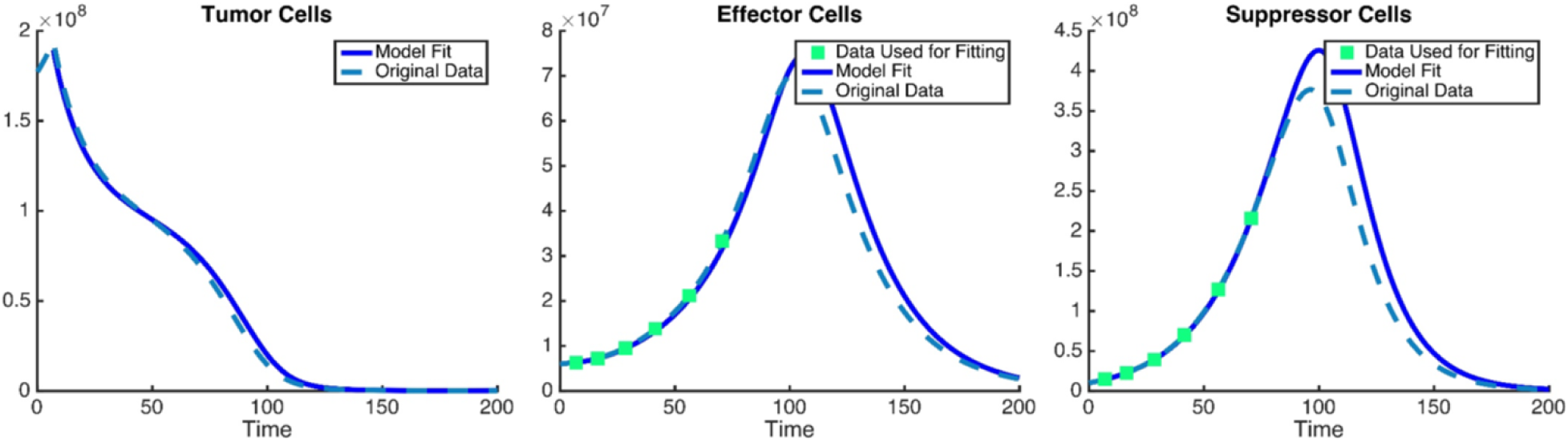
Parameter determination for a virtual patient treated with immunotherapy. Original virtual patient data is shown in a light blue dashed line, data points used for parameter estimation are shown in green squares, and parametrized model simulations in a solid blue line. We determine that six weekly data points were enough to recapture tumor behavior.

The final step is to obtain an optimal treatment strategy, which can be easily implemented in our model once all parameters are determined. Figure 5 shows tumor dynamics of a virtual patient undergoing various treatments, namely intermittent and continuous single drug treatment with mTORi, TKi, and NIVO. Intermittent mTORi treatment gives better prognosis than a continuous strategy for this specific patients. However, TKi works best with continuous administration, whereas NIVO shows no difference.

**Figure 5:**
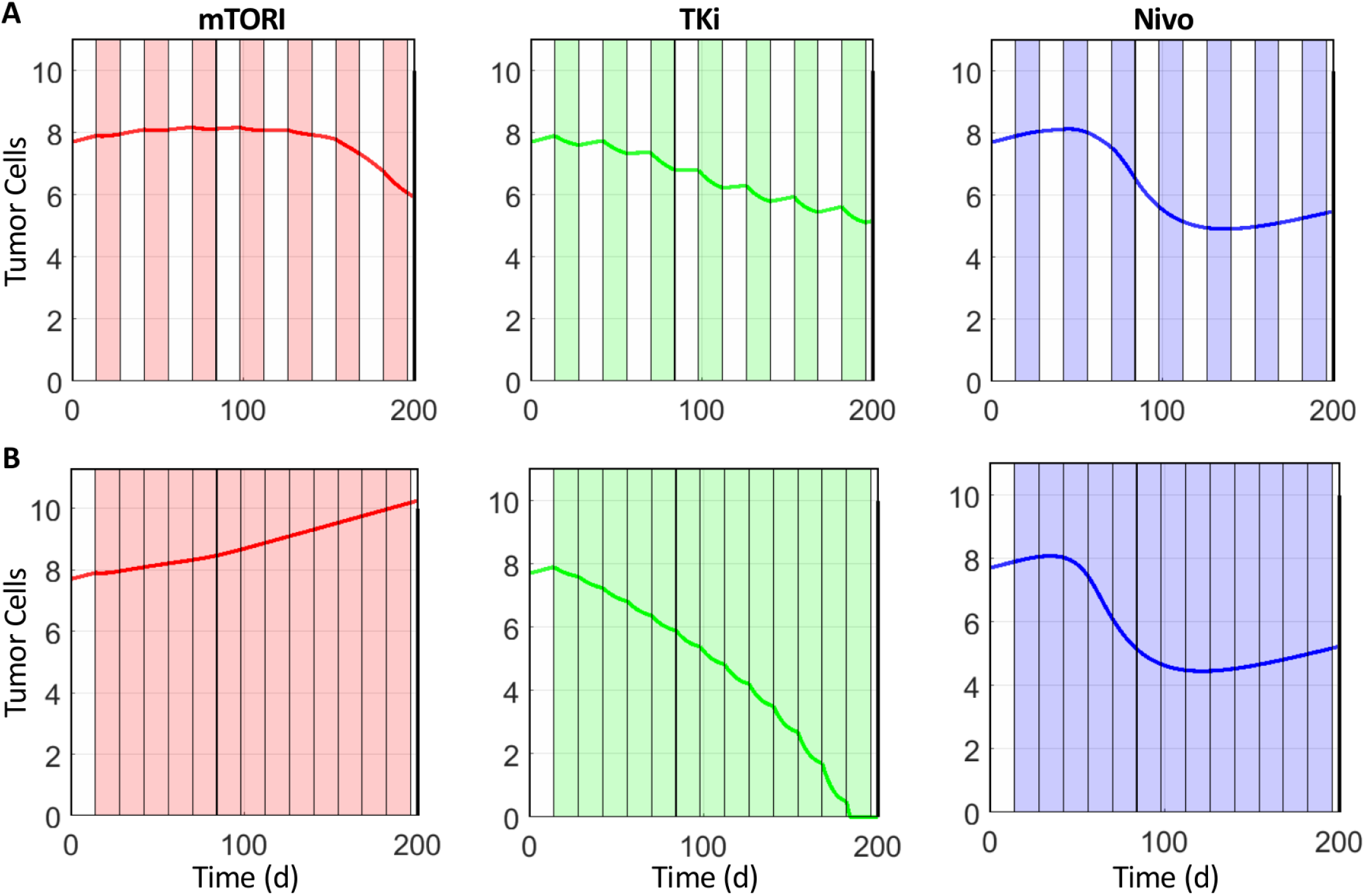
Comparison between continuous and intermittent treatment. Single-drug intermittent and continuous treatment were considered with mTORi, TKi, and Nivo. For this particular virtual patients, intermittent treatment was beneficial using mTORi, detrimental with TKi, and neutral using Nivo when compared to continuous treatment.

## 4. Discussion

RCC are highly heterogeneous, and with limited biomarkers to access heterogeneity clinicians may not be able to determine an individual patient’s prognosis accurately. Identifying individual patients tumor composition and personalizing treatment and treatment combinations has the potential to improve quality of life and increase survival. We propose that using imaging data and extracting immune markers via noninvasive blood tests could help initialize a mathematicl model for patient-specific treatment simulations. During treatment with either mTORi, TKi, or Nivo, weekly blood counts will be used to determine the sensitivity parameter associated with that treatment. Subsequently, the other single drug treatments will be given, specifying Equations (2) for the patient. We would then perform *in silico* simulations to determine the best treatment strategy for the patient. Similar to adaptive control approaches, treatment would begin, treatment response observed, model calibrated and response predicted, and repeated.

Patients with RCC show little to no response to traditional treatment strategies. Targeted therapies have been proposed as a better alternative by general guidelines. However, the diverse histology and genetic assembly make it difficult to find a treatment strategy that will benefit all patients. Evolutionary, personalized strategies can produce a better prognosis. Mathematical models can help clinicians choose such treatment. Our proposed model is capable of capturing tumor-immune interactions, as well as treatment mechanisms. In order to use such a model we propose NLR as a biomarker for treatment response. Using a reasonable number of blood samples, we can obtain enough NLR information to parameterize our model to recreate the tumor behavior of a specific patient. Optimal treatment can then be found by running *in silico* simulations.

## Acknowledgments

We will like to thank Dr. Alexander Anderson, for organizing the 8^*th*^ Annual Moffitt IMO workshop in evolutionary therapy, where this project was developed.

